# Rhes protein transits from neuron to neuron and facilitates mutant huntingtin spreading in the brain

**DOI:** 10.1101/2021.08.27.457956

**Authors:** Uri Nimrod Ramírez-Jarquín, Manish Sharma, Neelam Shahani, Yuqing Li, Siddaraju Boregowda, Srinivasa Subramaniam

## Abstract

Rhes (*RASD2*) is a thyroid hormone-induced gene that regulates striatal motor activity and promotes neurodegeneration in Huntington disease (HD) and tauopathy. Previously, we showed that Rhes moves between cultured striatal neurons and transports the HD protein, polyglutamine-expanded huntingtin (mHTT) via tunneling nanotube (TNT)-like membranous protrusions. However, similar intercellular Rhes transport has not yet been demonstrated in the intact brain. Here, we report that Rhes induces TNT-like protrusions in the striatal medium spiny neurons (MSNs) and transported between dopamine-1 receptor (D1R)-MSNs and D2R-MSNs of intact striatum and organotypic brain slices. Notably, mHTT is robustly transported within the striatum and from the striatum to the cortical areas in the brain, and Rhes deletion diminishes such transport. Moreover, we also found transport of Rhes to the cortical regions following restricted expression in the MSNs of the striatum. Thus, Rhes is a first striatum-enriched protein demonstrated to move and transport mHTT between neurons and brain regions, providing new insights on interneuronal protein transport in the brain.

## INTRODUCTION

The brain’s striatum plays a critical role in motor, cognitive and psychiatric functions, and its dysregulation results in neurological and neurodegenerative illnesses, such as Huntington disease (HD). The most common cell types in the striatum are the medium-sized spiny neurons (MSNs), and during postnatal brain development, the MSNs show the highest expression of Rhes (*RASD2*), a thyroid hormone-induced gene that regulates striatal motor activity (*1–3*). Rhes is enriched in the synaptic fractions, associates with membranes via the farnesylation domain, and mediates dopamine-related behaviors and G-protein coupled receptor and protein kinase A signaling (*4–7*). In response to psychostimulants, such as amphetamines or cocaine, Rhes also forms protein-protein complexes and alters proteomics in the striatum, while also inhibiting locomotor activities (*8, 9*). Rhes also regulates analgesia, tolerance, and dependency behaviors related to opioids [10].

In addition to its N-terminal GTPase domain, Rhes also possesses a C-terminal SUMO E3-like domain that promotes SUMOylation of diverse substrates including mHTT and promotes its solubility and toxicity in cell and animal HD models (*8, 10–19*). Rhes interacts with mammalian target of rapamycin (mTOR) kinase and promotes L-DOPA-induced dyskinesia in Parkinson disease (*20, 21*). Apart from its role in striatal diseases, Rhes is also linked to tau pathology, and its mislocalization in human neurons is considered a hallmark of tauopathies (*22, 23*). These results indicate that Rhes orchestrates neuronal abnormalities associated with neurodegenerative diseases, but the precise mechanisms of its action remain largely unknown. However, given the known importance of neuronal communication in brain function, cell-to-cell communication is likely to be important.

One intriguing new mode of cell communication occurs via transient and fragile membranous protrusions, such as tunneling nanotubes (TNTs) or cytonemes (henceforth TNT-like processes) (*24–33*). TNT-like processes are distinct from neurites or filopodia in cell culture in shape, size, length and strength. Currently, there are no cellular markers that distinguish TNT-like protrusion from neurites or filopodia. However, they are mostly F-actin positive, characteristically do not adhere to the substratum, extend up to 100 µm in length, physically connect two cells, and are vulnerable to common fixation methods (*34, 35*). These TNT-like processes are readily observed in various cell types in culture and in vivo and can be induced in cancer tissues (*36–40*). TNT-like processes can transport organelles, such as endosomes, lysosomes, and mitochondria, to neighboring cells (*41, 42*). Drosophila cytonemes regulate morphogenetic signals during development and can function as glutamatergic synapses (*43–45*). The emerging evidence indicates that TNT-like protrusions may play a critical role in key organism development and growth, but their roles and regulation in the brain remain unclear (*24, 46*).

We serendipitously discovered that Rhes promotes the formation of TNT-like membranous protrusions that transport cargoes such as endosomes and lysosomes (*46–48*). Interestingly, Rhes itself is transported via the TNT-like protrusions and can interact with lysosomes and mitochondria in the neighboring cells (*46–48*). Notably, in HD, Rhes causes a several-fold enhancement of the cell-to-cell transport of mHTT, indicating a cell non-autonomous role of Rhes in HD. Although the exact mechanisms by which cargoes of Rhes-mediated TNTs, such as mHTT, enter the acceptor cell are unknown, our live-cell time-lapse imaging studies have revealed the involvement of endocytic-like delivery mechanisms (*46–48*).

These data, taken together, indicate that the Rhes moves from cell to cell via the TNT-like membranous protrusions, but whether this movement and mHTT transport can occur in the intact brain is not known. Here, we address this question by developing new reporter tools and Cre recombinase transgenic mouse models and by live-cell imaging in organotypic brain slices and mouse brain that express MSN-specific reporter Rhes.

## RESULTS

### Rhes promotes protrusions and moves from D2R to D1R-MSN in cortico-striatal organotypic brain slices

We showed that Rhes travels between cells in the TNT-like protrusions in primary neurons and cell lines (*47–49*). But whether Rhes can induce TNT in MSNs of the striatum, where it is highly expressed in not known. Here we determined using live cell imaging if Rhes can induce TNT-like protrusion in primary MSNs and in a cytoarchitecturally intact 3D brain model. We established Flex (or “flip-excision”) genetic switch (Cre-On) replication-deficient adeno-associated virus (AAVs) with PHP.eB serotype, the most efficient vector that transduce neurons (*50*), encoding either EGFP or EGFP-Rhes. This approach restricts the reporter signal to Cre-expressing neurons; therefore, only if the reporter can move from neuron to neuron will the signal be found in non-Cre expressing neurons. We used EGFP as a reporter for Rhes because of its photostability, which allow live-cell monitoring of the dynamic membranous protrusions.

First, we determined if Rhes can induce TNT-like protrusions in primary D2R-MSNs prepared from D2R^Cre^ mice and infected with AAV Cre-On EGFP or AAV Cre-On EGFP-Rhes (Fig. 1A). We found that the D2R^Cre^-MSNs with AAV Cre-On EGFP expression showed typical neurites characteristic of morphologically crooked and bifurcated structures emanating from the cell body (Fig. 1B, inset *b*, blue arrow). In contrast, Cre-On EGFP-Rhes induced the formation of long (50-100 µm), straight, EGFP-Rhes positive TNT-like protrusions emanating from cell body (open arrow), as well as from the neurites (closed arrow) of the D2R MSNs (Fig. 1C, inset *c*, arrow). These Rhes-induced TNTs also contained characteristic round vesicle-like puncta at the surface (Fig. 1C, inset *c1*, arrowhead), consistent with prior observations in striatal neuronal cells (*47–49*).

**Figure 1.**
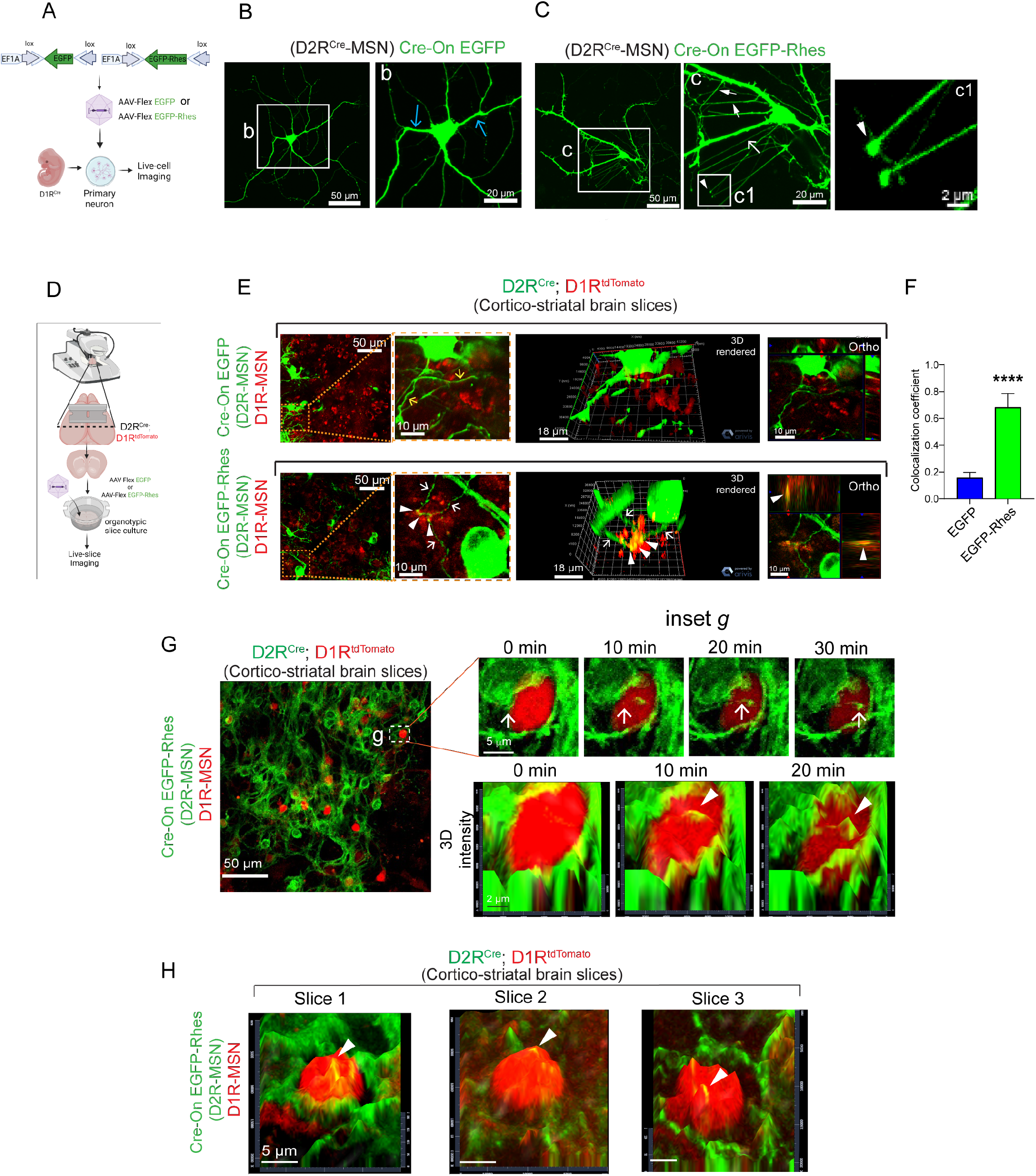
Rhes promotes TNT-like protrusions in MSNs. **(A)** AAV Flex Cre-On viral vector design and infection of AAV.PHP.eB Cre-On EGFP or EGFP-Rhes viral particles, into the D1R^Cre^ (*Drd1a*^*Cre*^) mice primary medium spiny neurons (MSNs). **(B, C)** Confocal images of 8-10 DIV D2R^Cre^-MSN expressing AAV Cre-On GFP **(B)** or AAV Cre-On GFP-Rhes **(C)** (MOI 10). Inset *b*, blue arrow indicates crooked and highly bifurcated neurites. Inset *c*, white open arrow indicates straight TNT-like protrusions from cell body of the GFP-Rhes expressing D2R^Cre^ neuron. Closed arrow indicates TNT-like protrusion from the neurites of the GFP-Rhes expressing D2R^Cre^ neuron. Arrowhead indicates GFP-Rhes positive vesicle-like blub commonly observed in Rhes-induced TNT-like protrusion. **(D)** Organotypic cortico-striatal brain slice culture from D2R^Cre^;D1R^tdTomato^ mice infected with AAV.PHP.eB Cre-On EGFP or EGFP-Rhes viral particles. **(E)** Confocal live-slice images and insets of organotypic cortico-striatal brain slices from D2R^Cre^;D1R^tdTomato^ mice transduced with Cre-On EGFP or Cre-On EGFP-Rhes AAV viral particles. Yellow and white arrow show neuronal processes of GFP or GFP-Rhes from Cre(+) D2R^Cre^ MSNs (green), respectively. The arrowhead shows GFP-Rhes puncta colocalization in Cre (–) D1R^tdTomato^ MSN (red). 3D rendering and orthogonal (ortho, single plane) display shows EGFP-Rhes puncta are inside the D1R^tdTomato^ MSN (arrowhead). **(F)** Pearson’s coefficient for colocalization between EGFP or EGFP-Rhes and tdTomato (n = 12, D1R^tdTomato^ neurons from 3 slices); data are mean ± SEM; Student’s t-test, ****p<0.0001. **(G)** Confocal time-lapse imaging of brain slices of D2R^Cre^-MSN (green) and D1R^tdTomato^ MSN (red) (D2R^Cre^;D1R^tdTomato^ mice) infected with AAV Cre-On EGFP-Rhes. Inset *g* (upper panel) shows EGFP-Rhes-positive TNT-like protrusions (0–30 mins) in the brain slices connecting D2R^Cre^-MSN (green) to D1R^tdTomato^-MSN (red). Inset *g* (lower panel) 3D intensity shows dynamic movement of Rhes dots from D2R^Cre^-MSN into D1R^tdTomato^ MSN (arrowhead) at different time points (0–20 min). **(H)** Confocal time-lapse imaging of brain slice showing 3D intensity of dynamic movement of Rhes dots from D2R^Cre^-MSN into D1R^tdTomato^ MSN (arrowhead) from three different brain slice experiments.

Other characteristic features distinguishing TNTs-like structures from neurites or filopodia in culture are their sensitivity to commonly used fixation and ability to stretch above the substratum. (*25, 51–53*). We found that Rhes-induced TNTs in D2R-MSN are disintegrated by fixative, paraformaldehyde (PFA) (sFig. 1A, B, dotted region, arrow), while certain protrusions that are most likely neurites remained intact (sFig. 1A, B, dotted region, arrowhead, compare DIC image). PFA treatment also destroyed Rhes-induced TNT-like protrusions (arrow) while sparing filopodia-like structures (arrowhead) in the striatal neuronal cell line (sFig. 1C, D). We also found that Rhes positive TNT-like protrusions are distinguishable from Rhes positive neurites (sFig. 1E). As shown in two examples, we found Rhes positive neurites are attached to the substratum (sFig. 1E, F, inset *e*-*f*, arrowhead). In contrast, Rhes positive TNT-like protrusions hover above the substratum and connect to nearby cell body (sFig. 1E, inset *e*, arrow) or neuronal processes (sFig. 1F, inset *f*, arrow).

Taken together, these results demonstrate that Rhes induces characteristic TNT-like structures that are distinct from neurites or filopodia in the primary MSNs. The data also show that these structures are highly fragile, further emphasizing the necessity of extracting dynamic information about TNT-like structures from live-cell imaging and reporter tools.

We next determined whether TNT-like structures are formed by Rhes in vivo. Two distinct types of MSNs, expressing either D1R or D2R, have been well characterized in the striatum. Approaches involving in situ anatomical studies and transgenic (Tg) reporter mice have now established that less than 5% of the MSNs co-express D1R and D2R, and these double positive MSNs are distributed uniformly throughout the striatum (*54–63*).

We used the organotypic cortico-striatal brain slices from postnatal 4-8-day old pups from D2R^Cre^;D1R^tdTomato^ mice and transduced the slices with AAV Cre-On EGFP or Cre-On EGFP-Rhes particles (Fig. 1D). As expected, the D2R neurons (Cre +) showed EGFP or EGFP-Rhes signals, marked by green fluorescence. Only a few EGFP-only positive protrusions (yellow arrow) were observed in the Cre-On EGFP cultures, but with no clear localization with D1R^tdTomato^ (red) neurons (Fig. 1E, upper panel). By contrast, we observed multiple EGFP-Rhes-positive protrusions (white arrow) interacting and colocalizing (arrowhead) with D1R^tdTomato^ (red) neurons (Fig. 1E, lower panel). A 3D rendering and orthogonal projections further confirmed a clear colocalization of EGFP-Rhes protrusions with D1R-MSNs (Fig. 1D, arrowhead, Fig. 1F). Unlike their appearance in dissociated cultures (Fig. 1B), the EGFP-Rhes positive TNT-like protrusions in slices appeared crooked, similar to those seen in mouse cornea explants of transgenic mice expressing Cx_3_cr1^GFP^, CD11c^eYFP^, or MHC class II^GFP^ (*64–66*). These results indicate that Rhes induces TNT-like protrusions in a complex interconnected 3D organotypic brain slices.

We also imaged EGFP-Rhes positive protrusions from the D2R^Cre^-MSNs traversing through the D1R^tdTomato^-MSNs by time-lapse confocal imaging of organotypic cortico-striatal brain slices (Fig. 1G, inset *g*). We found a time-dependent induction (zero to 30 minutes) of EGFP-Rhes positive protrusions from the D2R^Cre^-MSNs, which associates with the D1R^tdTomato^-MSNs (Fig. 1G, inset *g*, arrow, upper panel). Using 3D intensity, we showed that these protrusions merge with the D1R^tdTomato^-MSN surface (Fig. 1G, inset *g*, arrowhead, lower panel). This was consistently observed in three independent slice preparations, indicating potential Rhes delivery into D1R-MSN (Fig. 1H, arrowhead). Slices that express Cre-On EGFP alone did not show any obvious merging of EGFP protrusions with D1R-MSNs (sFig. 2).

**Figure 2.**
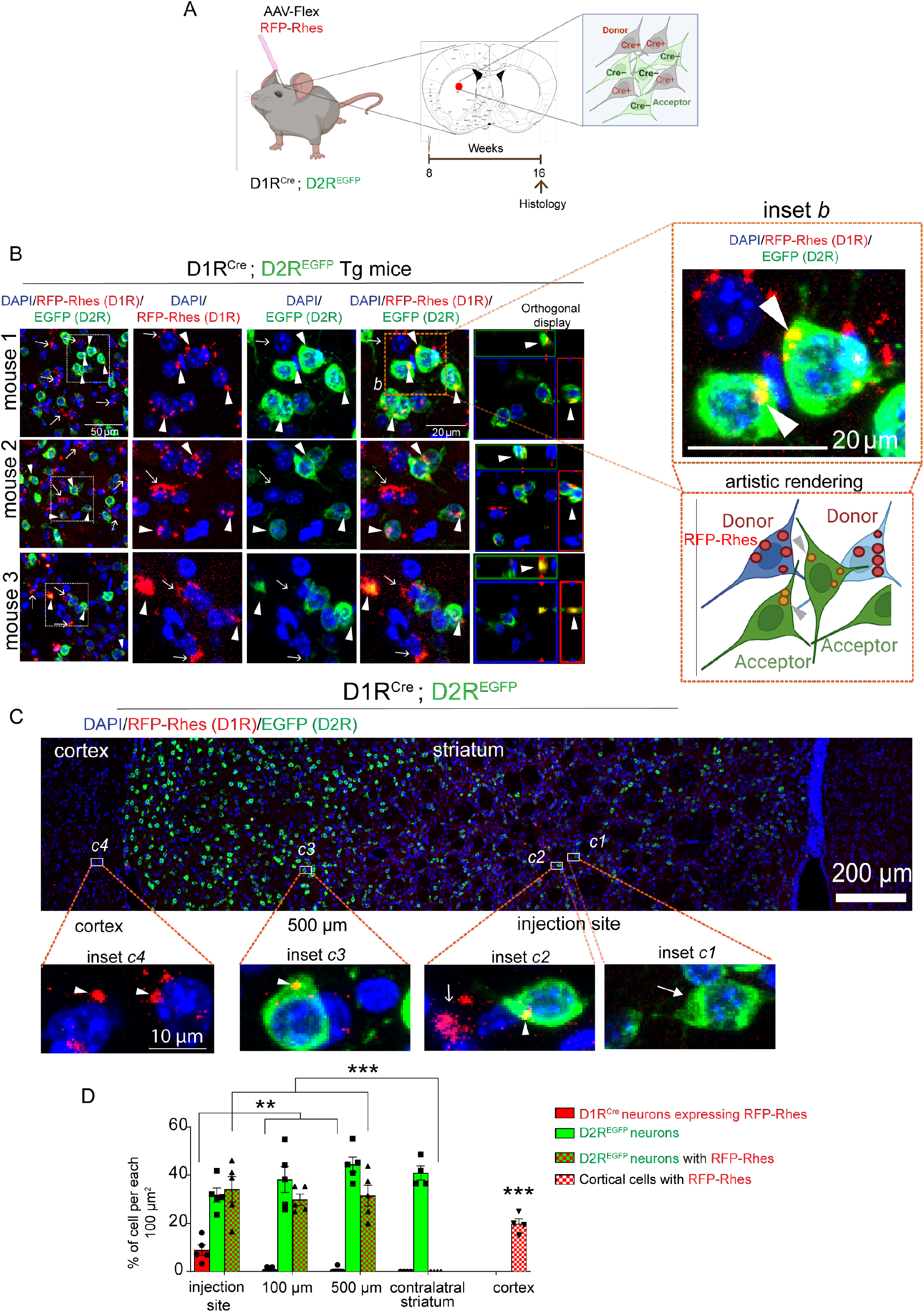
Rhes moves from D1R-MSN to D2R-MSN in vivo. (**A**) Cre-On in vivo model to investigate transport of Rhes from D1R-MSN to D2R-MSN in the striatum (using ML = ±1.6, AP = +1.0; DV = –3.6 coordinates) of D1R^Cre^;D2R^EGFP^ mice injected with AAV Cre-On RFP-Rhes. D1R-MSN will serve as Cre+ donor neuron and D2R-EGFP as Cre-acceptor neuron. (**B**) Confocal images of brain sections from three different D1R^Cre^;D2R^EGFP^ mice injected with Cre-On RFP-Rhes in the striatum. Arrow indicates expression of RFP-Rhes in D1R^Cre^(+) neurons. Arrowhead indicates RFP-Rhes expression in D2R^EGFP^ Cre (–) neurons. Arrowhead in the orthogonal display shows RFP-Rhes puncta are colocalized with D2R^EGFP^ neuron. Artistic rendering of inset b is shown. **(C)** A horizontal reconstruction of confocal images of D1R^Cre^;D2R^EGFP^ mice injected with Cre-On RFP-Rhes in the striatum. At the injected site, inset c1 shows D2R^EGFP^ neuron (closed arrow) and inset c2 shows D1R^Cre^ neurons expressing RFP-Rhes (open arrow) and D2R^EGFP^ neurons with RFP-Rhes (arrowhead). Inset c3 shows RFP-Rhes (arrowhead) in D2R^EGFP^ neuron, 500 µm away from the injected site and inset c4 shows RFP-Rhes in the ipsilateral cortical cells. (**D**) Bar graph shows quantification of the % of indicated neurons from the injected site, and 100 µm and 500 µm away from the injection in the striatum, ipsilateral cortex (as shown in C), as well as contralateral striatum. (n = 5/injection/mixed sex). Data are mean ± SEM, **p < 0.01, ***p < 0.001, Two-way ANOVA, Bonferroni post hoc test.

These results showed that Rhes can induce membranous protrusions and connect with the neighboring neurons in an intact 3D brain architecture.

### Rhes moves between D1R-MSN to D2R-MSN in the striatum

We next investigated the possible in vivo Rhes transport between MSNs by crossing D1R^Cre/+^ mice with D2R^EGFP/+^ mice. We tagged Rhes with TurboRFP, because of its intracellular stability, which allows RFP to be measured over a longer period of time. We confirmed that the AAV reporter construct expresses RFP-Rhes in a Cre-dependent manner and forms TNT-like protrusions (arrow) containing vesicular puncta-like bulb in their edge (arrowhead) in striatal neuronal cells in culture (sFig. 3A, B).

**Figure 3.**
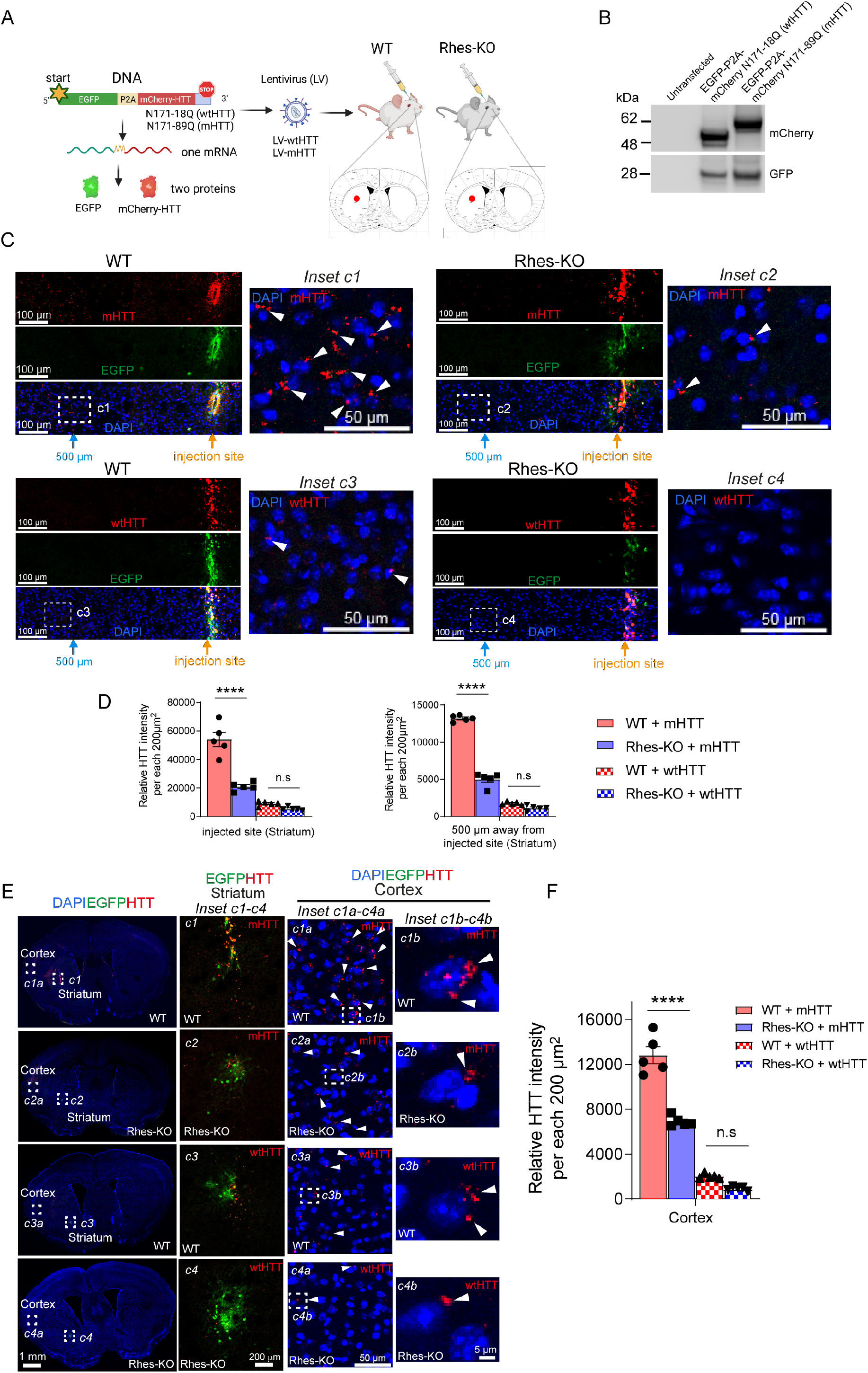
Rhes promotes cell to cell transport of mHTT in vivo. **(A)** Graphical representation of EGFP-P2A-mCherry N171-18Q (wtHTT) or EGFP-P2A-mCherry N171-89Q (mHTT) lentiviral vectors and injection of lentivirus into the striatum (using ML = ±1.6, AP = +1.0; DV = –3.6 coordinates) of WT or Rhes KO mice. **(B)** Western blotting of striatal neuronal cells expressing wtHTT or mHTT. **(C)** Confocal image of the horizontal reconstruction of a portion of brain section showing expression of EGFP (green) and wtHTT or mHTT (mCherry, red) at the injection site and 500 µm away from the injection site in WT and Rhes KO mice. Insets *c1-c4* show enlarged portion of a selected region in the 500 µm away from the injection site in the striatum. Arrowhead indicates perinuclear localization of mHTT. Cell nuclei stained with DAPI (blue). **(D)** Bar graph shows quantification of intensity of HTT in 200 µm^2^ striatal area in the injected site, and 500 µm away from the injected site (n = 5/injection/mixed sex). Data are mean ± SEM, ****p < 0.0001; n.s-not significant, One-way ANOVA, Tukey’s multiple comparisons test. **(E)** Confocal images of brain sections from WT and Rhes KO mice showing the expression of EGFP (green) and wtHTT or mHTT (mCherry, red) and cell nuclei stained with DAPI (blue). Insets *c1-c4* show magnified images from striatum and inset *c1a-c4a* and *c1b-c4b* from cortex. Arrowhead shows expression of mHTT or wtHTT in the cortex. **(F)** Bar graph shows quantification of intensity of HTT in 200 µm^2^ striatal area in the ipsilateral cortex. (n = 5injection/mixed sex). Data are mean ± SEM, ****p < 0.0001; n.s-not significant, One-way ANOVA, Tukey’s multiple comparisons test..

We then injected Cre-On AAV-RFP-Rhes or AAV-RFP (control) particles into the striatum in one hemisphere of 2-month-old D1R^Cre^;D2R^EGFP^ Tg mice brain, where D1R-MSN (Cre+) is considered as donor and D2R-EGFP (Cre-) as acceptor (Fig. 2A). At 8 weeks post-injection, the brain was perfused, and sections were processed for confocal imaging with DAPI as nuclear marker. RFP-Rhes (red, Fig. 2B, arrow) signals in the D1R^Cre^-MSN and EGFP (green) signals in D2R^EGFP^-MSN were detected in the striatum (Fig. 2B). In addition, we found a robust RFP-Rhes signal associated with D2R-EGFP^+^ neurons in the striatum, and this was consistent in three different mice (Fig. 2B, arrowhead). The RFP-Rhes signals were colocalized with EGFP neurons (Fig. 2B, orthogonal display, arrowhead) and were found in the perinuclear cytoplasm of the D2R MSNs (Fig. 2B, inset *b*, arrowhead). Using artistic rendering, we depicted the transported RFP-Rhes from D1R-MSN donor to D2R-MSN acceptor (Fig. 2B, inset *b*, lower panel).

We also determined the numbers of D1R (RFP-Rhes), D2R (EGFP), and D2R neurons containing RFP-Rhes in 100 µm^2^ sized striatal areas at the injection site, 100 µm and 500 µm away from the injection site and in the cortical areas of the brain sections of the D1R^Cre^;D2R^EGFP^ Tg mice (Fig. 2C, insets *c1-c3*, Fig. 2D).

At the injection site, we found 30% of D2R^EGFP^ only neurons (Fig. 2C, *inset c1* closed arrow, Fig. 2D) and 10% of D1R^Cre^ MSNs expressing RFP-Rhes (Fig. 2C, *inset c2* open arrow, Fig. 2D). We also observed at the injected site, ∼30% of D2R^EGFP^ MSNs positive for one or more RFP-Rhes puncta (Fig. 2C, *inset c2* arrowhead, Fig. 2D). Even at 100 µm and 500 µm away from the injection site, the 30% of D2R^EGFP^ neurons remained positive for RFP-Rhes (Fig. 2C, *inset c3* arrowhead, Fig. 2D), indicating that Rhes transits from D1R-MSNs to D2R-MSNs in the striatum. No similar colocalization was observed in the contralateral side of the striatum, indicating that Rhes is unable to move from ipsilateral striatum to contralateral striatum (Fig. 2D). Interestingly, we also observed strong signals for RFP-Rhes in the cortex of D1R^Cre^;D2R^EGFP^ mice (Fig. 2C, *insets c4* arrowhead, Fig. 2D), indicating Rhes movement from striatum to cortex.

Taken together, these results confirmed that Rhes can move from D1R-MSNs to D2R-MSNs within the striatum as well as from D1R-MSNs to the cortex in the mice brain.

### Rhes promotes mHTT spreading in the brain

We previously showed that Rhes is a major driver of the cell-to-cell transportation of mHTT in immortalized striatal cells (*48*). Our finding that Rhes is transported between neurons in vivo prompted us to investigate whether mHTT can also be transported by endogenous Rhes. We prepared lentiviral (LV) bicistronic vectors expressing EGFP and mCherry-tagged HTT containing N-terminal 171 aa with 18Q (LV-wtHTT) or 89Q (LV-mHTT) poly-glutamine (Q) repeats, separated by the P2A sequence (*67*) (Fig. 3A). In this scenario, EGFP and mCherry-HTT will be expressed as two separate proteins in an infected cell, thereby allowing the investigation of HTT transport by monitoring the mCherry fluorescence. We first validated the LV vectors and confirmed similar expression levels of two separate proteins by western blotting in striatal neuronal cell line (Fig. 3B). We then stereotaxically injected LV-wtHTT or LV-mHTT unilaterally into the striatum of 2-month-old WT and Rhes KO mice. After 2 months, the brains were fixed and processed for EGFP and mCherry fluorescence detection by confocal microscopy. As expected, we observed EGFP (green) and mCherry (HTT, red) co-expression at the injection site of the striatum (Fig. 3C). We also detected the mCherry-mHTT expression alone in the 200 µm^2^ area around the injection site, as well as 500 µm away from the injection site in the WT mice, indicating mHTT movement within the striatum (Fig. 3C, inset *c1*, arrowhead, Fig. 3D); however, the mCherry-HTT signal intensity was markedly reduced in the Rhes KO mice (Fig. 3C, inset *c2*, arrowhead, Fig. 3D). Moreover, little wtHTT was transported within the striatum compared to mHTT, and no significant differences were detected between WT and Rhes KO mice (Fig. 3C, *insets c3, c4*, Fig. 3D).

Remarkably, besides striatum (Fig. 3E, *inset c1-c4*), a strong mCherry-mHTT expression was observed in the cortex of the WT mice, which was much diminished in the Rhes KO mice cortex (Fig. 3E, inset *c1a-c4a*, arrowhead, Fig. 3F). When compared to mHTT, only few wtHTT puncta were seen in WT and it was not significantly different than Rhes KO cortex (Fig. 3E, arrowhead, Fig. 3F). Most mHTT and wtHTT puncta were distributed perinuclearly in the cortical cells of WT and Rhes-KO brain (Fig. 3E, inset *c1b-c4b*, arrowhead).

These observations indicate that the mHTT protein is transported between neurons and to cortical areas and that Rhes is a major driver for such transport.

### Rhes moves from the striatum to the cortical areas of the brain

The above data (Figs. 2 and 3) suggested that Rhes can move to the cortical region, indicating propensity for extrastriatal migration. First, we examined whether Rhes might be transferred between cortical neurons in vitro. We co-cultured cortical neurons of EGFP Tg mice and *CamKII*^*Cre*^ Tg mice and infected them with AAV Cre-On RFP or AAV Cre-On RFP-Rhes viral particles (Fig. 4A). Three days after the infection, we observed two distinct neuronal populations expressing Cre-On RFP in CamKII^Cre^ neurons (donor) and EGFP neurons (acceptor) without any apparent overlap (Fig. 4B). Little or no RFP signal was found in EGFP neurons (Fig. 4B, Inset *b*). Remarkably, a robust transport of Cre-On RFP-Rhes from CamKII^Cre^ (donor) neuron was found in the perinuclear regions of the EGFP (Cre –) cortical acceptor neurons (Fig. 4B, inset *b1*, arrowhead). Orthogonal rendering further confirmed that RFP-Rhes was colocalized with the EGFP neurons (Fig. 4B, inset *b1*, arrow). Overall, RFP-Rhes but not RFP alone is transported efficiently between cortical neurons (Fig. 4C).

**Figure 4:**
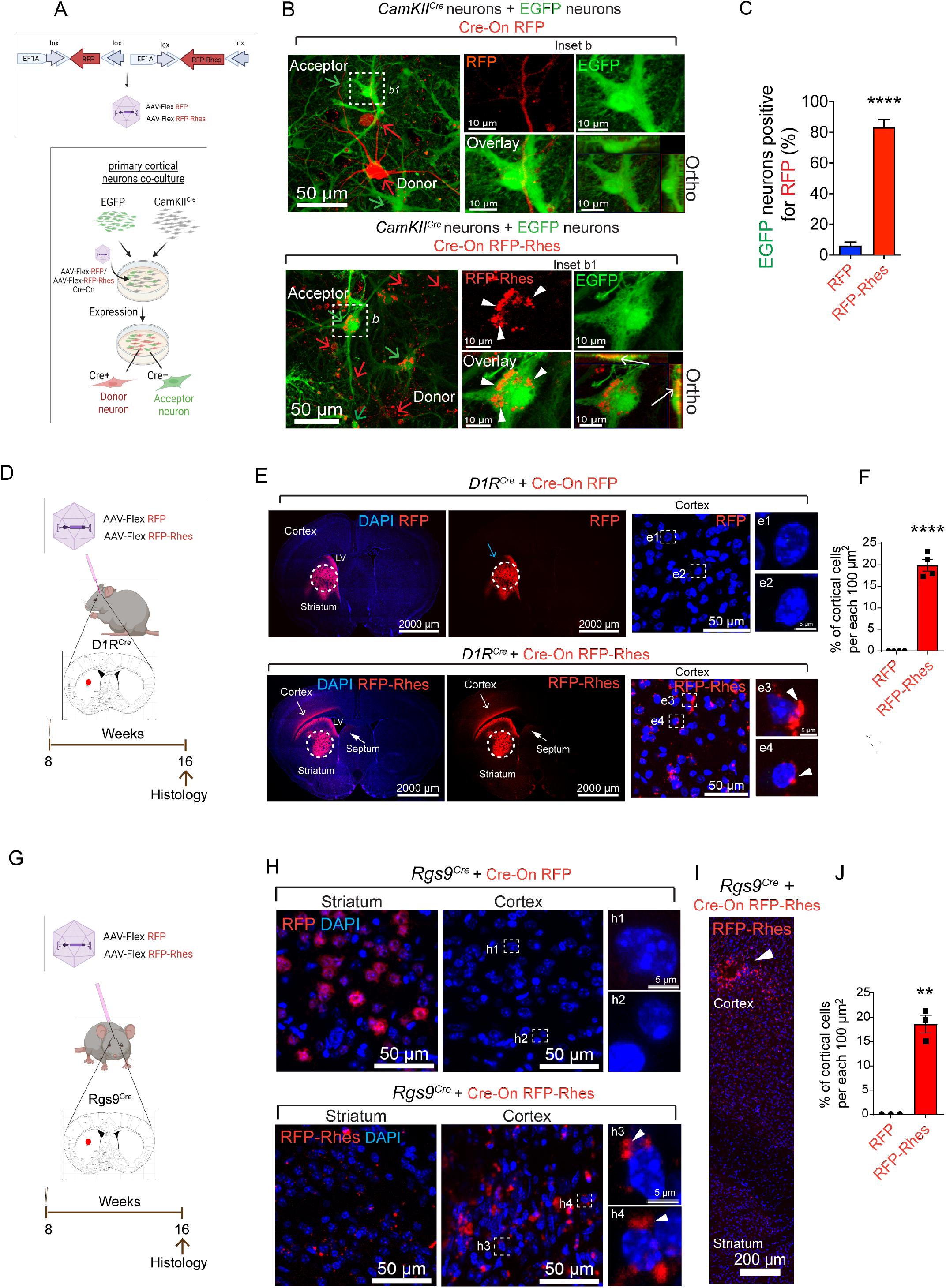
Rhes moves from striatum to cortex. (**A**) Cre-On AAV Flex viral vector design (upper panel) and primary cortical neuron co-culture model (lower panel) to investigate Rhes transport in vitro. Primary cortical neurons were co-cultured from Cam-KII^Cre^ mice (donor neuron) and EGFP Tg mice (acceptor neuron) and infected with Cre-On AAV Flex RFP or Cre-On AAV Flex RFP-Rhes. (**B**) Confocal images of primary cortical neuron co-culture from Cam-KII^Cre^ mice and EGFP Tg mice. Red arrow indicates Cam-KII^Cre^ neurons (Cre+ donor neurons) expressing Cre-On RFP alone (upper panel) or Cre-On RFP-Rhes (lower panel). Green arrow indicates EGFP neurons (Cre–, acceptor neuron). Inset *b and b1*, enlarged portion of EGFP acceptor neuron. Arrowhead indicates Cre-On RFP-Rhes in EGFP acceptor neuron. Orthogonal display show colocalization of Cre-On RFP-Rhes in the EGFP acceptor neuron (arrow). (**C**) Quantification of EGFP neurons positive for RFP signal (%) for RFP Cre-On (n= 29) and RFP-Rhes Cre-On groups (n = 41). Data are mean ± SEM; Student’s t-test, ****p<0.0001. **(D)** Stereotaxic injection of AAV Cre-On RFP or RFP-Rhes viral particles using ML = ±1.6, AP = +1.0; DV = –3.6 coordinates into the D1R^Cre^ (*Drd1a*^*Cre*^) mice striatum. **(E)** Coronal brain section of the D1R^Cre^ mice injected with Cre-On RFP or RFP-Rhes in the striatum with cell nuclei stained with DAPI (blue) and dotted circle showing RFP or RFP-Rhes signals (red) in the injected area. RFP alone signal in the striatum (blue arrow). RFP-Rhes signals in the cortical region (open arrow) and septal regions (closed arrow). Insets *e1-e4* shows the high magnification image of the cortical region. Insets *e3-e4* shows perinuclear cytoplasmic accumulation of Rhes in the cortical cells (arrowhead). (**F**) Quantification of percent of cortical cells in the cortex of D1R^Cre^ mice injected with Cre-On RFP or RFP-Rhes in the striatum showing the expression of RFP/RFP-Rhes. Data are mean ± SEM; Student’s t-test, **p <0.01 (n = 4/injection). (**G**) Stereotaxic injection of AAV Cre-On RFP or RFP-Rhes viral particles into the Rgs9^Cre^ mice striatum using ML = ±1.6, AP = +1.0; DV = –3.6 coordinates. (**H**) Confocal image of striatum and cortical regions of Rgs9^Cre^ mice injected with Cre-On RFP or RFP-Rhes in the striatum with cell nuclei stained with DAPI (blue). Insets *h1-h4* shows the high magnification image of the cortical region. Insets *h3-h4* shows perinuclear cytoplasmic accumulation of Rhes in the cortical cells (arrowhead). (**I**) A vertical reconstruction of confocal images of Rgs9^Cre^ mice brain injected with RFP-Rhes in the striatum. Arrowhead indicates an accumulation of RFP-Rhes in the cortex. (**J**) Quantification of percent of cortical cells expressing RFP-Rhes in Rgs9^Cre^ mice injected with Cre-On RFP-Rhes in the striatum. Data are mean ± SEM; Student’s t-test, **p <0.01 (n = 3/injection).

Next, we set out to determine further if the intraregional transport of Rhes can occur from the striatum to cortex using D1R^Cre^ mice (Fig. 4D). We stereotaxically injected AAV Cre-On RFP (control) or Cre-On RFP-Rhes particles unilaterally into the adult mouse striatum of *Drd1a*^Cre/+^ Tg mice (Fig. 4D), in which D1R-positive MSNs are primarily restricted to the striatum, as confirmed in *Drd1a*^*EGFP*^ mice (sFig. 4A). At ∼8 weeks after injection, the *Drd1a*^Cre/+^ Tg mouse brains were perfused, and sections were analyzed for RFP and RFP-Rhes signal by confocal microscopy. As expected, we found Cre-On RFP expression highly restricted to the striatum (Fig. 4E, *upper panel*, blue arrow). By contrast, the Cre-On RFP-Rhes signals were found in the striatum, but strong signals were also observed in the cortical areas (layer V and VI) of the brain (open arrow), with some signal also evident in the septum region (closed arrow) (Fig. 4E, *lower panel*). The RFP-Rhes, but not the RFP signal alone, was consistently found as perinuclear cytoplasmic structures in the cortical cells of the *Drd1a*^Cre/+^ mouse cortex (Fig. 4E, insets *e1-e4*, arrowhead, and Fig 4F).

We further confirmed the cortical transport of Rhes in *Rgs9*^*Cre*^ mice (Fig. 4G) that primarily express Cre in the striatal MSNs (*68, 69*). As expected, striatal injection of Cre-On RFP in the *Rgs9*^*Cre*^ mouse striatum resulted in RFP expression effectively restricted to the striatum (sFig. 4B, Fig. 4H, *upper panel*). By contrast, striatal injection of Cre-On RFP-Rhes resulted in RFP-Rhes signals in the striatum and also cortical areas of the *Rgs9*^*Cre*^ mice (Fig. 4H, *lower panel*, & Figs. 4I, 4J). Signal from RFP-Rhes, but not RFP alone, was observed in the perinuclear cytoplasm of the cortical cells (Fig. 4H, insets *h1-h4*, arrowhead, and Fig 4J), consistent with the observations in D1R^Cre^ mice (Fig. 4E).

Taken together, these results indicated that Rhes protein can transit efficiently in vitro between primary cortical neurons and in vivo from the striatum to the cortex

## DISCUSSION

In this report, we provide compelling evidence that the brain-enriched protein Rhes can move from neuron to neuron in dissociated neuron preparations, in brain slices, and in intact mouse brains. Previous work has shown that prion proteins, as well as misfolded proteins, such as α-synuclein, amyloid, and tau, can move from neuron to neuron, although the mechanisms are not fully understood (*70, 71*). The Arc protein, which shows virus-like properties, can be transported between neurons via extracellular vesicles/exosomes (*72, 73*). Rhes does not share homology with prions or with Arc protein, nor is it present in exosomes (sFig. 5). The findings presented here broaden the transport mechanisms for structurally unrelated proteins between neurons.

**Figure 5.**
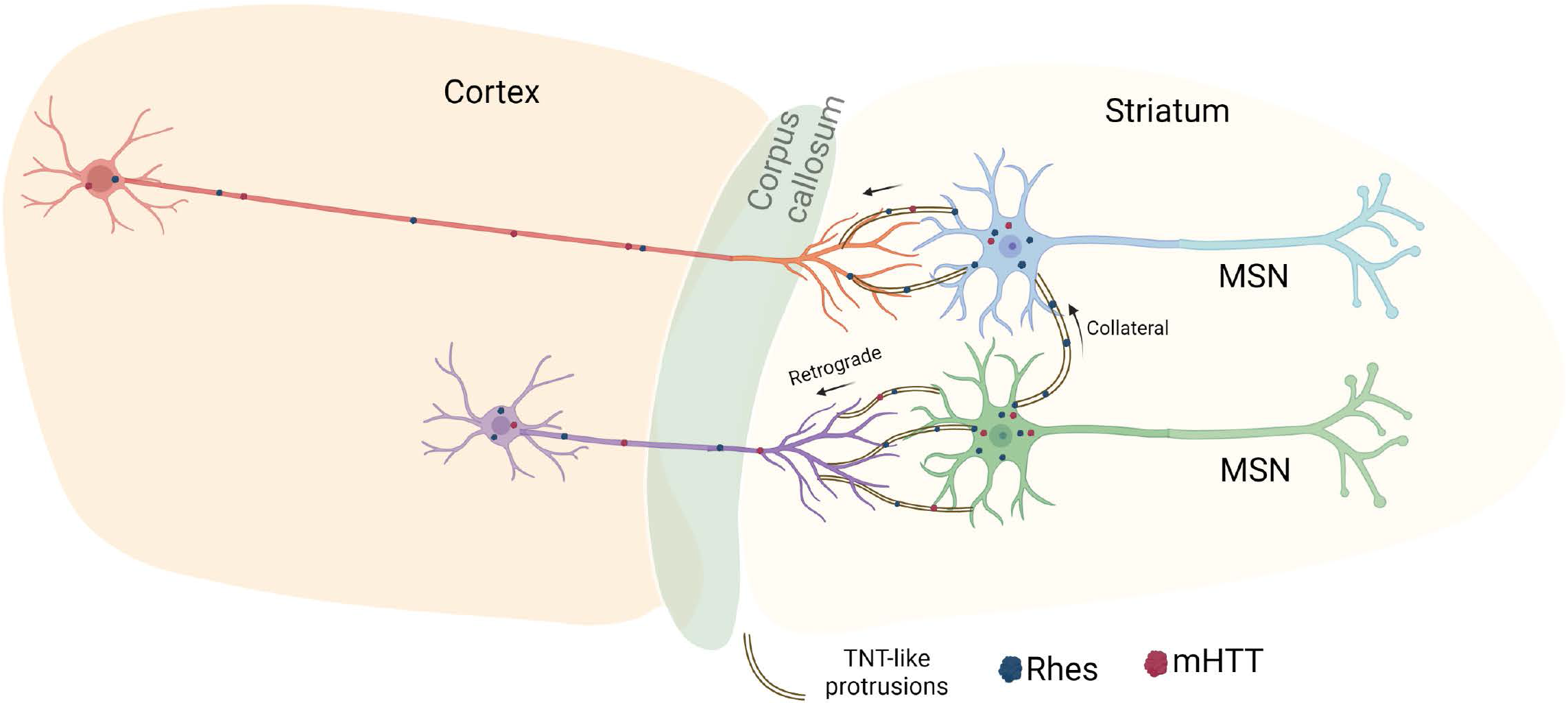
Neuron to neuron Rhes transport model. Our data indicate that Rhes moves between MSNs in the striatum as well as to the cortical areas. Live-cell imaging data from primary neurons and organotypic slices data indicate TNT-like membranous protrusions are the key routes Rhes contact the neighboring neurons. Thus, we predict that Rhes transports and facilitates mHTT movements in vivo potentially via the direct physical contact of neurons via membranous protrusions. Both collateral contact between MSNs and cortical-striatal contacts of MSN to the cortical projections may occur via TNT-like membranous protrusions.

Our data suggest that neuron-to-neuron Rhes transport involves TNT-like mechanisms. Rhes is expressed in PC12 cells (*10, 74*), in which TNTs were first reported (*75*). We confirmed that the depletion of endogenous Rhes in the PC12 cells diminishes TNTs, indicating Rhes is required for TNTs formation or maintenance (sFig. 6). Rhes, in addition to engineering actin-based TNT-like membranous protrusions, travels by this route and is delivered to the acceptor cells (*46–48*). Clues by which this occurs emerged from time-lapse cell imaging studies showing that the TNT-like protrusions touch the acceptor cells and deliver vesicle-like puncta in a process that resembles endocytosis-like mechanisms of cargo uptake (*46, 48*). The live-cell imaging of brain slices further demonstrated that the protrusions and their related delivery mechanisms are likely operational in vivo (Fig. 1) and that Rhes transport from neuron to neuron in the adult brain (Fig. 2) can occur by a protrusion-based mechanism (*46, 48*).

Anatomical and electrophysiological studies have shown that D1R and D2R MSNs are interconnected by collateral axons (*76–79*). Our present work showed that Rhes moves between MSNs, adding a new dimension to neuronal communication in the striatum. Beyond the MSNs, the interneurons of the striatum are also positive for EGFP-positivity in D2R^EGFP^ mice (*80*). Therefore, the possibility that Rhes might be transported to other cell types, such as interneurons in the striatum, cannot be ruled out.

Neuron to neuron communication via synapses and neuron to glia via extracellular fluid is necessary for neuronal functioning. Rhes-mediated TNT-like communication may provide a network of direct routes connecting the cell body and the synapses of neurons in the brain. Rhes is highly abundant in the neuronal synapses, where it may bridge synapses via the tiny TNT-like protrusions to transport proteins or vesicular cargoes depending upon the functional and metabolic demands of neurons and their disease state.

Previous studies have reported the cell-to-cell transport of mHTT in human HD patient striatum, as well as in mouse and Drosophila HD models (*81–86*). Extracellular vesicles and endocytosis are suspected to be involved in the mHTT delivery and uptake (*87, 88*), however, the details of the molecular mechanism that drives mHTT transport between neurons in the brain are unclear. Consistent with earlier work (*89*), we showed that mHTT can be transported by TNT-like protrusions and that Rhes accelerates this process several fold (*48*). In this report, we confirm that mHTT transport occurs in WT mice; however, the transport is markedly diminished in the Rhes-KO brain (Fig. 3), indicating that Rhes is a physiological mediator of mHTT transport in vivo.

Several studies have indicated that mHTT has prion-like properties that may contribute to its cell-to-cell transmission (*83, 86, 90–92*). Previous work showed that TNTs may serve as a route for the transport of prion-like protein between cultured cells (*93, 94*). An interesting possibility is that Rhes may increase the transmission of prion-like mHTT in vivo via the TNT-like processes. Recent work has shown that tau pathology and spreading can occur via TNTs, and Rhes is a critical determinant of tau pathology (*22, 23, 93, 95, 96*). Remarkably, the spread of tau pathology can occur from the striatum, and tau is also associated with HD pathology (*97–101*). We found Rhes interacts with striatal low-density lipoprotein (LDL) receptor-related protein 1 (LRP1) (*8*), which is involved in tau uptake and spread (*102*). Thus, Rhes may have a binary function in neurodegenerative disease pathology by promoting transport of both misfolded tau and mHTT from the striatum.

One intriguing question is how Rhes moves from the striatum to the cortex (Fig. 4). Our data supports the model that Rhes promotes the formation of TNT-like processes from the MSN to connect to the axonal projections arising from the cortex (corticostriatal fibers), thereby allowing retrograde transport of Rhes (Fig. 5). Retrograde transport of proteins, and particularly the plant protein horseradish peroxidase (HRP), is well documented. HRP is transported in membrane-bound vesicles within the axons of ganglion cells in the optic tectum of the chick and rat sciatic nerves (*103–105*). HRP injected into the striatum was later found in various regions of the cortex in macaque monkeys (*106*). Similar to HRP (*105*), Rhes is found localized with lysosomes in the acceptor cells (*48*), indicating that HRP and Rhes might share similar destination mechanisms. On lysosomes Rhes might influence mTORC1 signaling and autophagy (*107*). Besides lysosomes, Rhes is also localized to endosomes and damaged mitochondria in the acceptor cells (*47, 48*). Therefore, we predict that conserved intercellular retrograde transport and docking mechanisms are involved in transporting Rhes from the TNT-like protrusions of the striatal MSNs to the cortex (Fig. 5).

Our in vitro studies also indicated an involvement of Rhes in cell-to-cell transport of receptors, such as D1R, D2R, and histamine-3 receptor, but not of TMEM214, an ER-associated transmembrane protein (sFig. 7), suggesting that this specific transport of cargoes by Rhes may modulate cell autonomous signaling between neurons in an unprecedented manner. Future work should clarify the cellular and molecular mechanisms by which Rhes moves and transports cargoes between neurons and their physiological role in the brain.

In summary, our results demonstrate that Rhes transits and transports mHTT in the brain most likely involving protrusion-based communication routes between neurons and interconnecting neural pathways. The findings presented here indicate the potential of developing new therapeutic approaches for interfering with Rhes-mediated mHTT spreading in the brain to slow or prevent HD.

## MATERIALS AND METHODS

### Animals

Surgery and stereotaxic injections were made in adult animals (8 weeks old). Animals were housed in groups of three to five on a 12:12 h light-dark cycle and food and water were ad libitum provided. All protocols were approved by the Institutional Animal Care and Use Committee at The Scripps Research Institute, Florida.

For Rhes_Flex Cre-On expression we used previously well-characterized Cre or reporter transgenic mice. D1R^Cre^ mice were obtained from MMRRC (030989-UCD-Hemi-F; B6.FVB(Cg)-Tg(Drd1-Cre)EY262Gsat/Mmucd), D2R^Cre^ mice were purchased from MMRRC (032108-UCD-Hemi-F; B6.FVB(Cg)-Tg(Drd2-Cre)ER44Gsat/Mmucd), D2R^EGFP^ mice were purchased from MMRRC (000230-UNC-Hemi-M; Tg(Drd2-EGFP)S118Gsat/Mmnc). CamKII^Cre^ transgenic mice (005359 B6.Cg-Tg(Camk2a-cre)T29-1Stl/J, homozygous), D1R^Td-Tomato^ mice (016204; B6.Cg-Tg(Drd1a-tdTomato)6Calak/J, Hemi-M) and EGFP transgenic mice (006567; C57BL/6-Tg(CAG-EGFP)131Osb/LeySopJ, homozygous) were obtained from The Jackson Laboratory. RGS9^Cre^ mice were produced as described before (*68*). Double transgenic mice (D1R^Td-Tomato^/D2R ^Cre^, and D1R^Cre^/D2R^EGFP^) were obtained by crossing the male D1R^Td-Tomato^ with female D2R^Cre^ and the female D1R^Cre^ with male D2R^EGFP^, respectively. Female with the Cre transgenes were used for breading, according with the suggestions from the company. Heterozygous condition for the Cre transgene was used for all the experiments, with exception of RGS9-Cre, which was used in homozygous form.

#### Constructs details

Cre-On RFP and Cre-On Turbo RFP-Rhes was cloned in pAAV (FLEX Cre-On) vector under EF1A promoter (VectorBuilder). Similarly, the Cre-On EGFP and Cre-On EGFP-Rhes is cloned in pAAV (FLEX Cre-On) under CAG promoter (VectorBuilder). Human *RASD2* (NM_014310.3) sequences was used for Rhes expression. The details of vector and cloning can be found in sFig. 8. The vectors were packaged in AAV-PHP.eB serotype and ultra-purified using VectorBuilder service (www.vectorbuilder.com). The mCherry-HTT N171 18Q or 89Q were cloned in 3rd generation Lentiviral vector for bi-cistronic expression of EGFP and the gene of interest (Addgene, 24129) and transfected in HEK 293T cells along with packaging plasmids. After 24 hr. of transfection media was changed and cells were incubated for additional 48 hrs. Supernatant was collected and virus was purified, and the titer was determined by infecting HEK 293T cells by a serial dilution. pEGFP-N-Drd1 plasmid was a gift from Kirk Mykytyn (Addgene, 104358), GFP-DRD2 plasmid was a gift from Jean-Michel Arrang (Addgene, 24099), pH3R-mCherry-N1 plasmid was a gift from Dorus Gadella (Addgene, 84327), and Tmem214 pmRFP-N2 plasmid was a gift from Eric Schirmer (Addgene, 62047). Scrambled gRNA and rat-specific gRNA CRISPR constructs were custom made from VectorBuilder.

### Stereotaxic striatal injections

Intrastriatal surgery for AAV/lentivirus infusion was carried out using the stereotaxic coordinates as described in our previous studies (*11, 108*). Briefly, adult (8-10 weeks old) male and female mice were injected with the virus according to the designed experiment. For surgery the mice anesthetized through the constant delivery of isoflurane in oxygen while mounted in a stereotaxic frame (David Kopf Instruments). Unilateral injection was made into the striatum at the following coordinates: ML = ±1.6, AP = +1.0; DV = –3.6 from bregma. Viruses were injected in 0.5 µl volume. The following viruses were used: AAV RFP-Rhes (Flex) Cre-On 1.24 × 10^12^ gc/ml; AAV RFP (Flex) Cre-On 2.39 × 10^12^ gc/ml; AAV GFP-Rhes (Flex) Cre-On 1.28 × 10^12^ gc/ml; AAV GFP (Flex) Cre-On 2.12 × 10^12^ gc/ml; Lentivirus GFP-P2A-mCherry HTT N171-18Q 1.0 × 10^12^ gc/ml; Lentivirus GFP-P2A-mCherry HTT N171-89Q 1.0 × 10^12^ gc/ml.

### Tissue fixation for imaging

Tissue was minimally processed to avoid loss of endogenous fluorescent signal. Briefly, 8-weeks after the stereotaxic injection, mice were anesthetized and perfused with cold saline solution (10 ml, 1.5 ml/min rate perfusion), followed by 4% paraformaldehyde (10 ml, 1.5 ml/min rate perfusion). Mouse brains were collected and postfixed overnight in 4% paraformaldehyde, cryoprotected in a sucrose/PBS gradient at 4ºC (10, 20, 30%), and then embedded in Tissue-Tek OCT compound (Sakura). Coronal sections (20 µm) were collected on Superfrost/Plus slides, counterstained with DAPI, and mounted using Fluoromount-G mounting medium (ThermoFisher Scientific). Images were obtained with the Zeiss LSM 880 microscope and processed using ZEN software (Zeiss).

### Organotypic cultures

Organotypic cortico-striatal slices were prepared from transgenic postnatal 4-8 days old mouse pups of both sexes from D2R^Cre+/-^;D1R^tdTomato+/-^ mice, as described in our earlier work (*109*). In brief, the animals were quickly decapitated, and the brain removed, the cerebellum and frontal hemisphere were cut. Cortico-striatal coronal slices (350 µm thick) were obtained using a vibratome (Leica). Slices were collected and kept in dissection media [Minimum Essential media (MEM, M7278, Millipore Sigma), 1% GlutaMAX (ThermoFisher Scientific) and 1% penicillin/streptomycin (ThermoFisher Scientific)]. Single slice was placed on interface-style Millicell culture inserts (PICM0RG50, 30 µm, hydrophilic PTFE, 0.4 µm pore size) in 6-well culture plates containing 1 ml of sterile slice culture medium [50 % MEM, 25 % Basal medium eagle (BME, B1522, Millipore Sigma), 25% heat inactivated horse serum (ThermoFisher Scientific), 0.6 % Glucose (G8769, Millipore Sigma), 2 mM GlutaMAX, 1% antibiotic antimycotic (15240096, ThermoFisher Scientific). Brain slices were incubated at 37°C in 5% CO_2_. Two days later 800 µl culture medium was removed and replaced with 800 µl culture medium. On day 5 culture medium was replaced with low-serum Neurobasal-N1 medium (94.5% Neurobasal plus medium, 0.5% heat-inactivated horse serum, 1X N1 supplement, 2 mM glutamine, 0.6% glucose, 1% antibiotic antimycotic). Culture medium (Neurobasal-N1 medium) was exchanged every 3-to-4-days. On day 7 in vitro slice cultures were infected with 0.5 µl of Cre-On GFP or Cre-On GFP-Rhes using a droplet method (*109*). For live imaging (7-10 days after infection), slices were cut out of the insert (still attached to membrane) and were transferred into glass-bottom dishes (D11140H, Matsunami Glass) with a drop of imaging media (A14291DJ, ThermoFisher Scientific) and were imaged using a Zeiss LSM 880 microscope, and the videos were analyzed by using the ZEN software.

### Primary neuron culture, and co-culture experiments

Primary neuron culture was performed as described in Sharma and Subramaniam, JCB, 2019 (*48*). In brief, striata of postnatal day 1 D2R^Cre^ mice were removed and digested at 37°C for 15 min in a final concentration of 0.25% papain and resuspended in neuronal plating media (Neurobasal-A media; Thermo Fisher Scientific), with 5% FBS, 0.5 mM glutamax, and 1% penicillin-streptomycin. Tissues were dissociated by trituration with a pipette. Further, cells were plated in 35-mm glass-bottom dishes (D11140H; Matsunami) coated with 100 µg/ml poly-D-lysine at the density of 2 × 10^5^ cells per dish. Dishes were maintained in a 37°C, 5% CO2 incubator. After the cells adhered (1–3 h after plating), plating media were replaced with growth media (Neurobasal-A media, 2% B27, 0.5 mM glutamax, and 1% penicillin-streptomycin). Neurons were infected with AAV Cre-On EGFP or Cre-On EGFP-Rhes at DIV 10 (MOI 10). Confocal mages were acquired after 48 hr. in live cell imaging solution as described earlier (*48*). For fixation experiments, similar strategy was used for primary neuron culture as above. Live cell imaging was acquired of primary neurons or striatal neuronal cells expressing GFP-Rhes. Later cells were fixed with 1% PFA for 10 seconds and images were captured from the same field to assess the effect of PFA fixation on TNTs stability. For co-culture experiments, primary cortical neurons were prepared from CamK-II^Cre^ mice and EGFP Tg mice. Cells were plated in 1:1 ratio and cultured and infected using AAV Cre-On RFP or Cre-On RFP-Rhes (MOI 10) as mentioned above. Percentage of EGFP neurons positive for RFP signal for RFP Cre-On and RFP-Rhes Cre-On groups were quantified. PC12 stably expressing scramble or rat-Rasd2 gRNA were grown as described before (*107*).

### Confocal imaging

Fixed tissue and organotypic cultures were imaged using Zeiss LSM880 confocal system. Whole coronal reconstructions were acquired using a 20x objective, with a Z-stack of three planes. High magnification images and partial cortico-striatal reconstructions were acquired using a 63x objective, optical Zoom was adjusted according to the field of interest. Live imaging of organotypic slices were acquired using a 40x objective with a 0.5 optical zoom. Primary neurons, and striatal neuronal cells transfected with reporter Rhes and reporter cargoes were live-cell imaged as described before (*48*).

### Quantification

Confocal images were used to quantify the protein expression using the ImageJ software. Average from two to three sections from each mouse were used for group analysis, and average of two to three different areas from each region (injection site, at 100 µm and 500 µm from the injection site, contralateral striatum) were analyzed. Proportion of D1, D2 and cortical cells with Rhes, as well as D2 alone without Rhes, after injection was made by quantifying each kind of neuron in 100 µm^2^ areas from each region. Total number of nuclei stained with DAPI was consider as 100% for each 100 µm^2^ analyzed. Similarly, ipsilateral cortical areas intensity of RFP or RFP-Rhes were determined in D1R^Cre^ and Rgs9^Cre^ mice brain sections in 100 µm^2^ area.

Transportation of the mCherry-HTT (N171-18Q) protein or the mCherry-mHTT (N171-89Q) was measured using the ImageJ software, by quantifying the relative intensity for the HTT or mHTT expression (mCherry, red channel). Average of two areas (200 µm^2^) from each region (injection site, 500 µm from the injection site and cortex) were analyzed for group results (site injection was identified by the EGFP expression).

### Exosome isolation

The exosomes isolation was performed using a protocol described previously (*110, 111*). Briefly, the striatum tissue was dissected out, weighed, and transferred to a 50 ml tube containing 75 U/ml of collagenase in PBS (total volume of PBS was 6 ml). The tissue was then incubated in a shaking water bath at 37°C for a total of 30 min. Tube was mixed 3 times during incubation by gentle inversion (after 10 min incubation). The collagenase was diluted by adding ice cold PBS (up to 44 ml to make the total volume 50 ml). Once the tissue was settled down, around 48 ml PBS was discarded. The washing step was repeated one more time to further dilute out collagenase. Protease and phosphatase inhibitor (Millipore Sigma) were added to a final concentration 1× in remaining 2 ml PBS. Tissue was introduced to gentle pipetting (10 stroke). The dissociated tissue was spun at 300 × g for 5 min at 4°C, the pellet was collected for input and the supernatant was transferred to a fresh tube, spun at 2000 × g for 10 min at 4°C. Pellet was discarded and supernatant was spun at 10,000 × g for 30 min at 4°C. The supernatant was overlaid on sucrose cushion (10%, 20%, 30%, 40% and 50%). The gradient was spun for 3 h at 180,000 × g (average) at 4°C (SW41 Beckman ultra-centrifuge). After the spin the top of the gradient was removed and discarded, and the 10% fraction (F1) was designated number 1. Fractions 2 (20%), 3 (30%) 4 (40%) and 5 (50%) were subsequently collected and protein was precipitated via methanol chloroform method (*47*). Exosome isolation was confirmed by flotillin antibody in western blot.

#### Western blotting

Striatal neuronal cells (STHdh^Q7/Q7^) grown in growth medium containing Dulbecco’s modified Eagle’s medium with high glucose (Thermo Fisher Scientific) with10% FBS, transfected with indicated plasmids or infected with AAVs and lysed in RIPA buffer and equal proteins were loaded onto the SDS page and transferred as described before (*112*). Primary antibodies against-Rhes (1:500, FabGennix #RHES-101AP), GFP (1:5000, Cell Signaling, #2956), mCherry (1:5000, Novus Biologicals, #NBP2-25157), Flotillin (1:1000, Cell Signaling, # 18634) and secondary antibodies conjugated to HRP (1:10000, Jackson labs) were used.

### Statistical analysis

Data are presented as mean ± SEM as indicated. Variance was found to be similar between the groups tested. Statistical analysis was performed with a Student’s *t* test or one-way analysis of variance (ANOVA) followed by Tukey’s multiple comparison test or two-way ANOVA followed by Bonferroni post-hoc test. Significance was set at *p* < 0.05. All statistical tests were performed using Prism 9.0 (GraphPad software).

## ACKNOWLEDGEMENT

We would like to Dr. David Fitzpatrick, Dr. Ronald Davis and Dr. Baoji Xu for critical reading of the manuscript and for their constructive suggestions. We thank Melissa Benilous for her administrative help and the members of the lab for their continuous support and collaborative atmosphere. We like to thank members at the Max Planck Institute of Neuroscience, Jupiter, FL microscopy core for their help and expertise. This research was supported from NIH/NINDS R01-NS087019-01A1, NIH/NINDS R01-NS094577-01A1, a grant from Cure for Huntington Disease Research Initiative (CHDI) foundation and Scripps bridge funding. We also thank Biorender application tools for model preparation as described in the manuscript.

## Author contributions

S.S conceptualized the project and co-designed experiments with U.N.R.J. who performed mouse experiments, immunohistochemistry, immunostaining, confocal, colocalization analysis and analyzed the data. M.S designed IRES constructs, performed primary neuron experiments, and N.S co-designed and prepared brain slice experiments and in vivo injections as well as western blotting. Y. L generated the Rgs9^Cre^ mice. S.B interpreted the data and provided intellectual contribution. S.S wrote the manuscript with inputs from coauthors.

## LEGENDS

**Supplementary Figure (sFig) 1.**
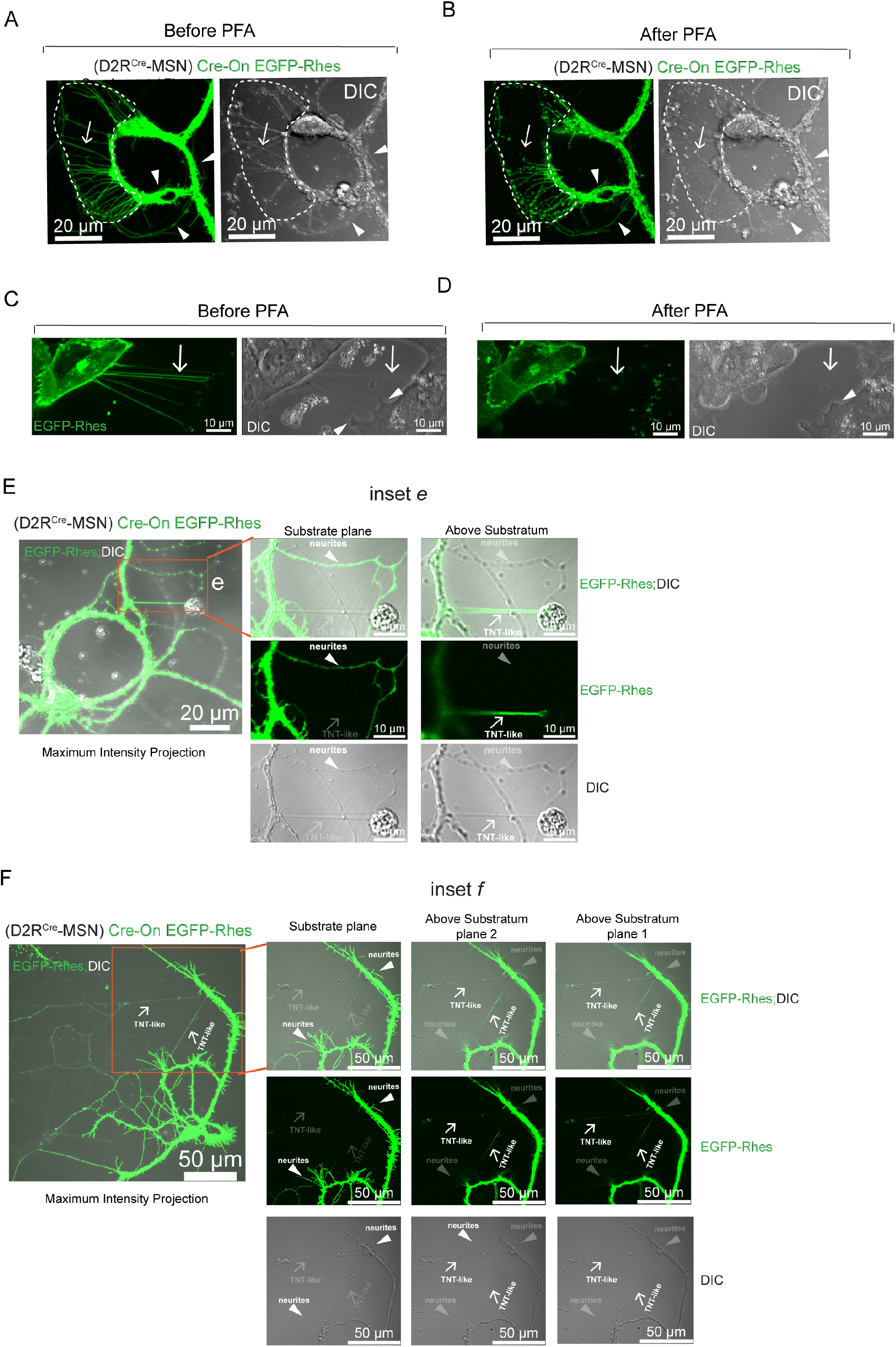
**(A-B)** Live-cell confocal imaging showing Cre-On EGFP-Rhes-induced TNT-like protrusion in D2R^Cre^-MSN before paraformaldehyde (PFA, 1%) fixation (*A*) and after PFA fixation (*B*). **(C-D)** Live-cell confocal imaging showing EGFP-Rhes-induced TNT-like protrusions in striatal neuronal cell lines before PFA fixation (*C*) and after PFA fixation (*D*). Differential interference contrast (DIC). Rhes-induced TNT-like protrusions (arrow) are destroyed after PFA treatment Arrowhead indicates neurite (A-B), filopodia-like protrusion (C-D) remain intact after PFA. **(E)** Confocal and DIC image of D2R^Cre^-MSN expressing EGFP-Rhes shows two different planes (substrate plane or above substrate). Inset *e*, arrowhead points to neurites visible in the substrate plane. At this plane the TNT-like cellular protrusion becomes out of focus (dim). Inset *e*, arrow points to the visible TNT-like protrusion above the substratum, where the neurites become out of focus (dim). **(F)** Confocal and DIC image of D2R^Cre^-MSN expressing Cre-On GFP-Rhes shows two different planes (substrate plane or above substrate). Inset *f*, arrowhead points to neurites visible in the substrate plane. At this plane, the TNT-like cellular protrusions become dim. Inset *f*, arrow points to the visible TNT-like protrusions above the substratum (plane 1 and plane 2). At these planes the neurite-like processes are dim.

**Supplementary Figure 2.**
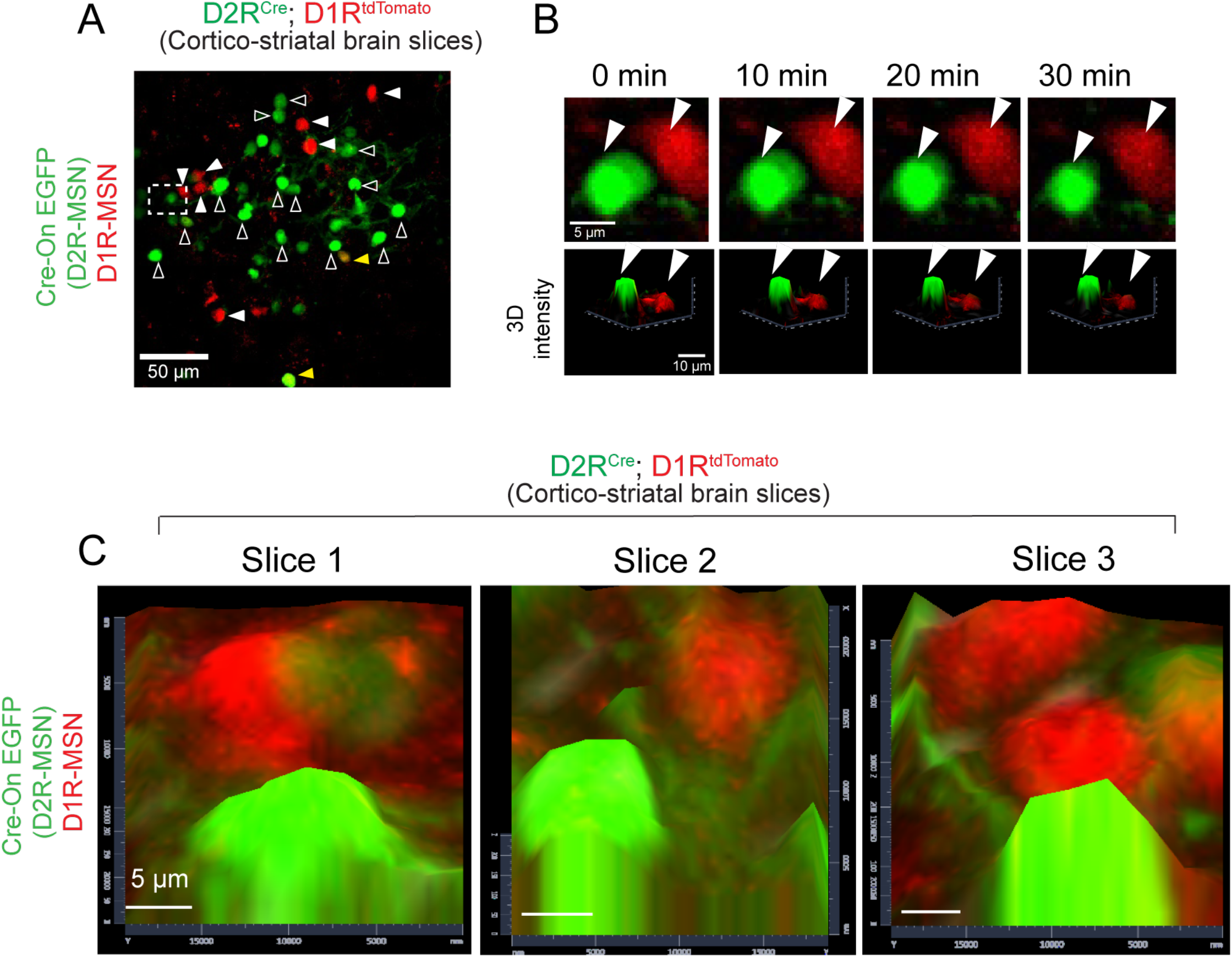
**(A)** Confocal time-lapse imaging of D2R^Cre^-MSN (green) and D1R^tdTomato^-MSN (red) from D2R^Cre^; D1R^tdTomato^ organotypic cortico-striatal brain slices infected with AAV Cre-On GFP **(B)** Inset (upper panel) shows GFP and tdTomato MSNs (arrowhead) time lapse imaging from 0–30 mins. 3D intensity inset (lower panel) shows lack of apparent association at different time points (0–30 min). **(C)** Confocal time-lapse imaging of the organotypic cortico-striatal brain slices infected with AAV Cre-On GFP showing 3D intensity of D2R^Cre^-MSN and D1R^tdTomato^-MSN from three different brain slice experiments.

**Supplementary Figure 3.**
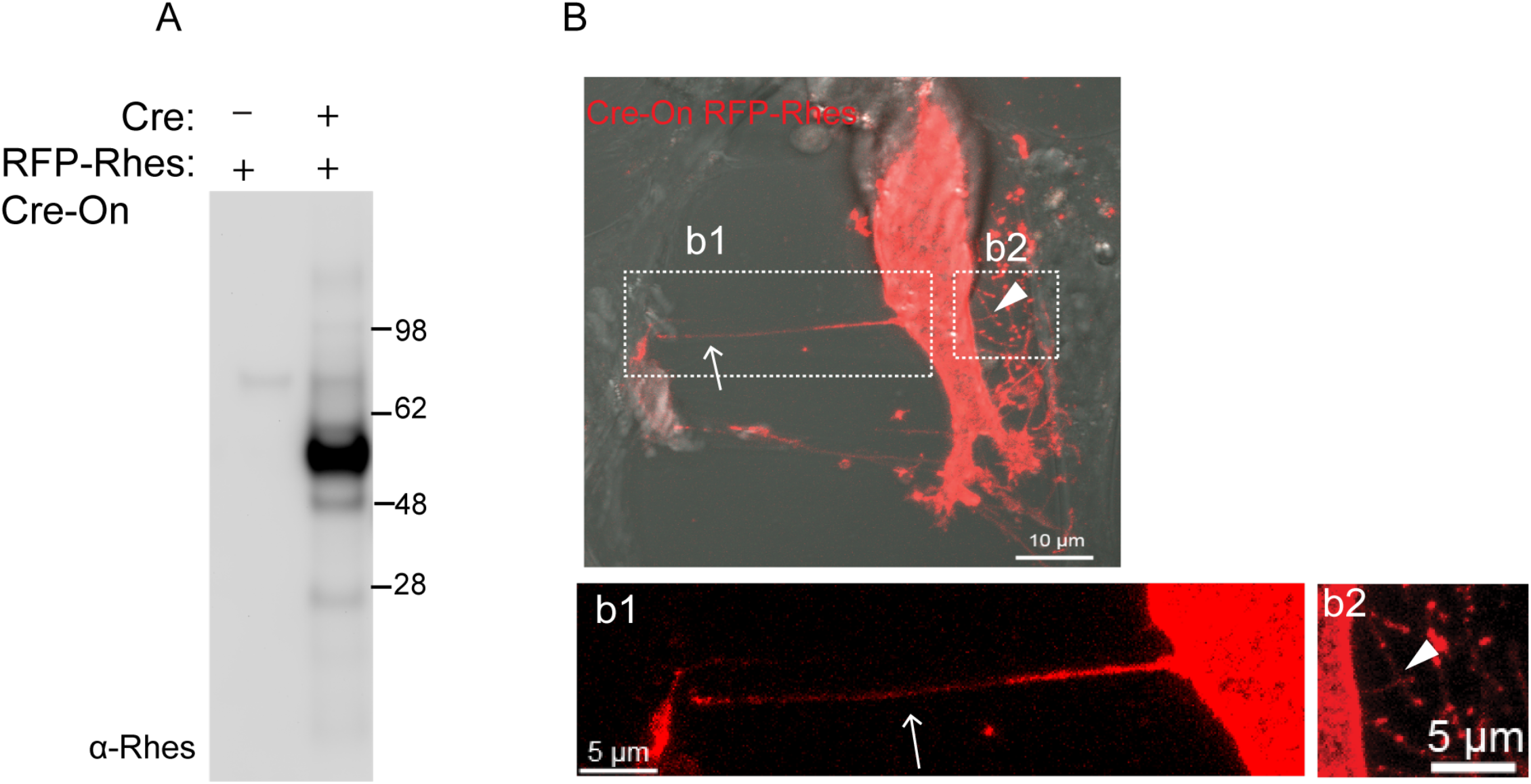
**(A)** Western blot for Rhes in the lysates from striatal neuronal cells transfected with or without cDNA of Cre and RFP-Rhes Cre-On vector. **(B)** Striatal cells expressing Cre-On RFP-Rhes and forming a long (*inset b1*, arrow) and short (*inset b2*, arrowhead) TNT-like cellular protrusions.

**Supplementary Figure 4.**
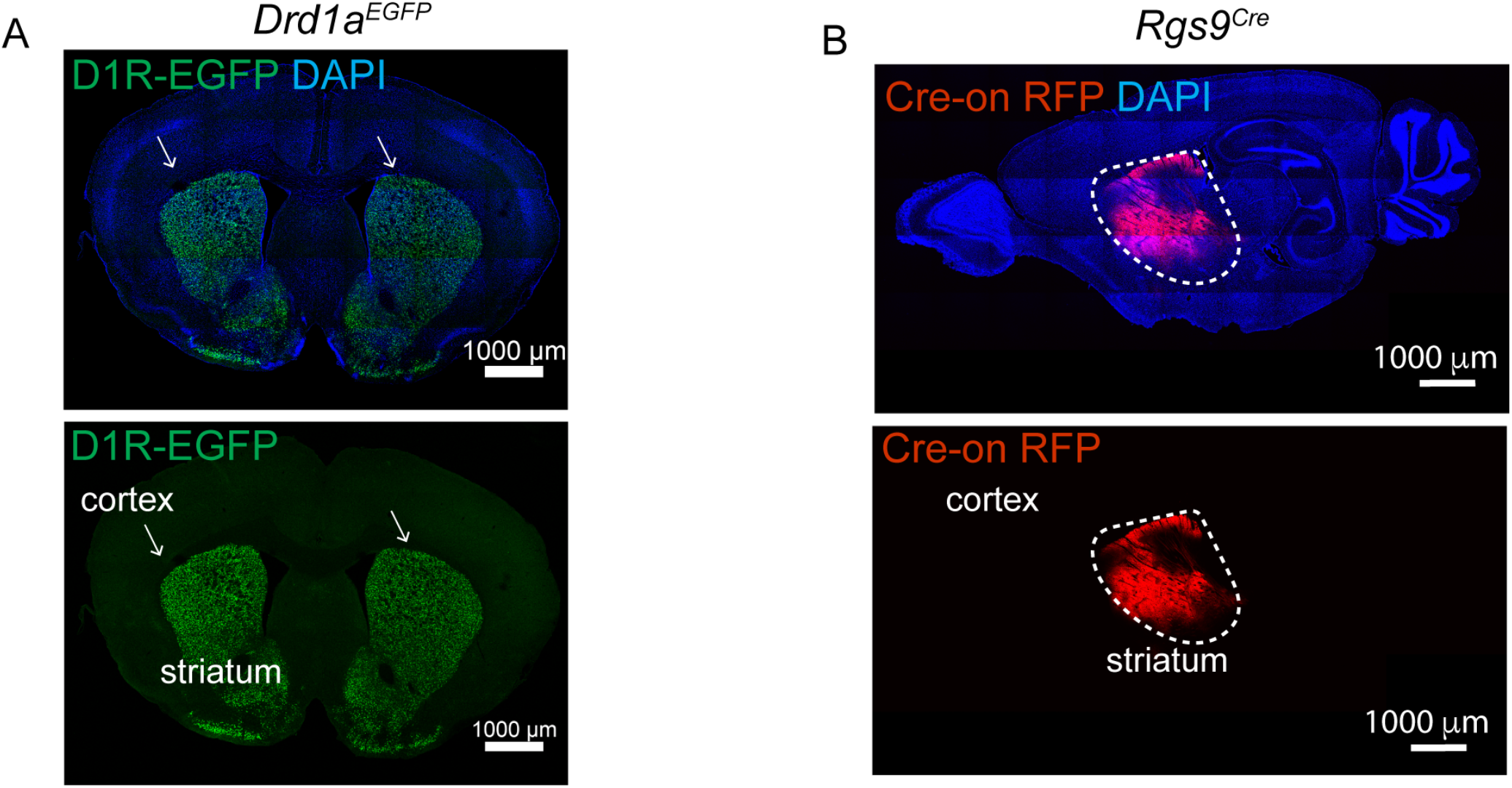
(**A)** Coronal brain sections of *Drd1a*^*EGFP*^ mice showing expression of EGFP in D1R-MSNs with nuclear stain (DAPI). **(B)** Sagittal brain section of *Rgs9*^*Cre*^ mice injected with Cre-On RFP in the striatum.

**Supplementary Figure 5.**
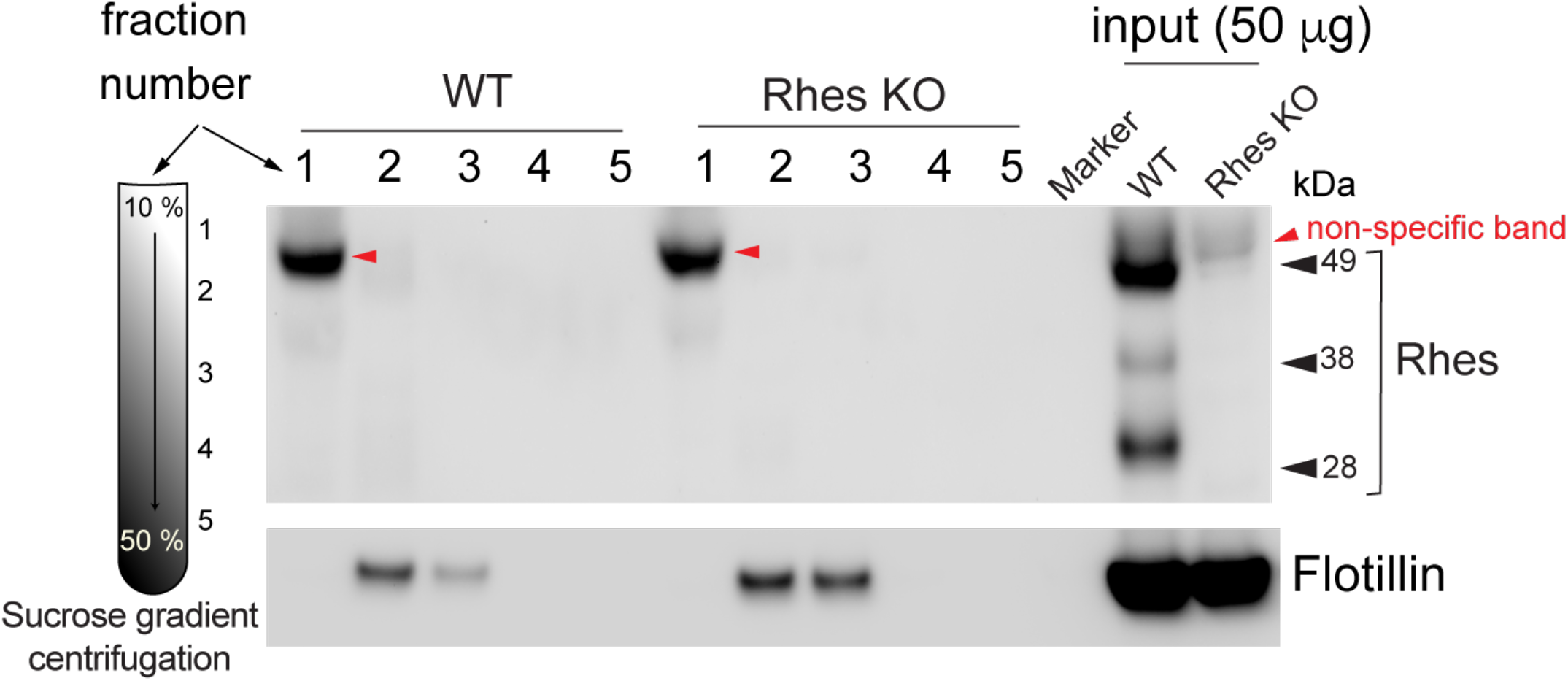
Western blot for Rhes and flotillin, an exosome marker, in the striatal lysate from WT or Rhes KO mice after differential sucrose gradient fractionation.

**Supplementary Figure 6.**
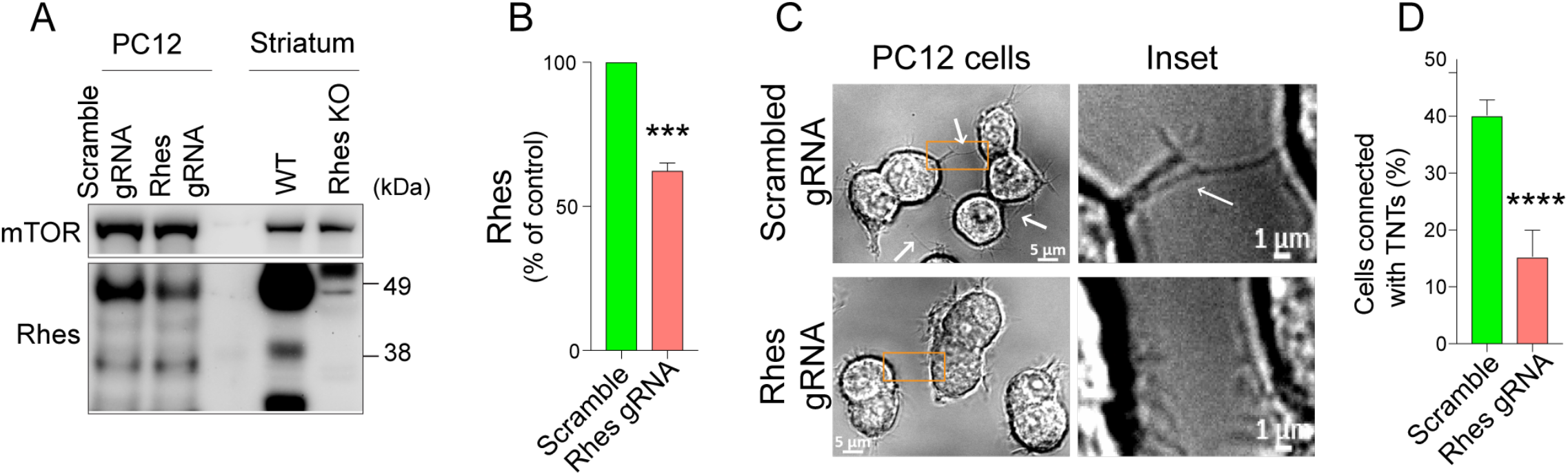
Rhes promotes TNTs in PC12 cells. **(A)** Western blot and quantification **(B)** of knock down of Rhes by CRISPR Rhes gRNA in PC12 cells that are stably selected with puromycin. WT and *Rhes*^*–/–*^ mice striatum used as positive control. **(C)** DIC live-cell imaging of stable PC12 cells and magnified insets. Arrows indicate TNTs. **(D)** % of cells connected with TNTs (scrambled n = 132; Rhes gRNA n =108). Mean ± SEM, unpaired t-test, ***p<0.001; ****p<0.0001.

**Supplementary Figure 7.**
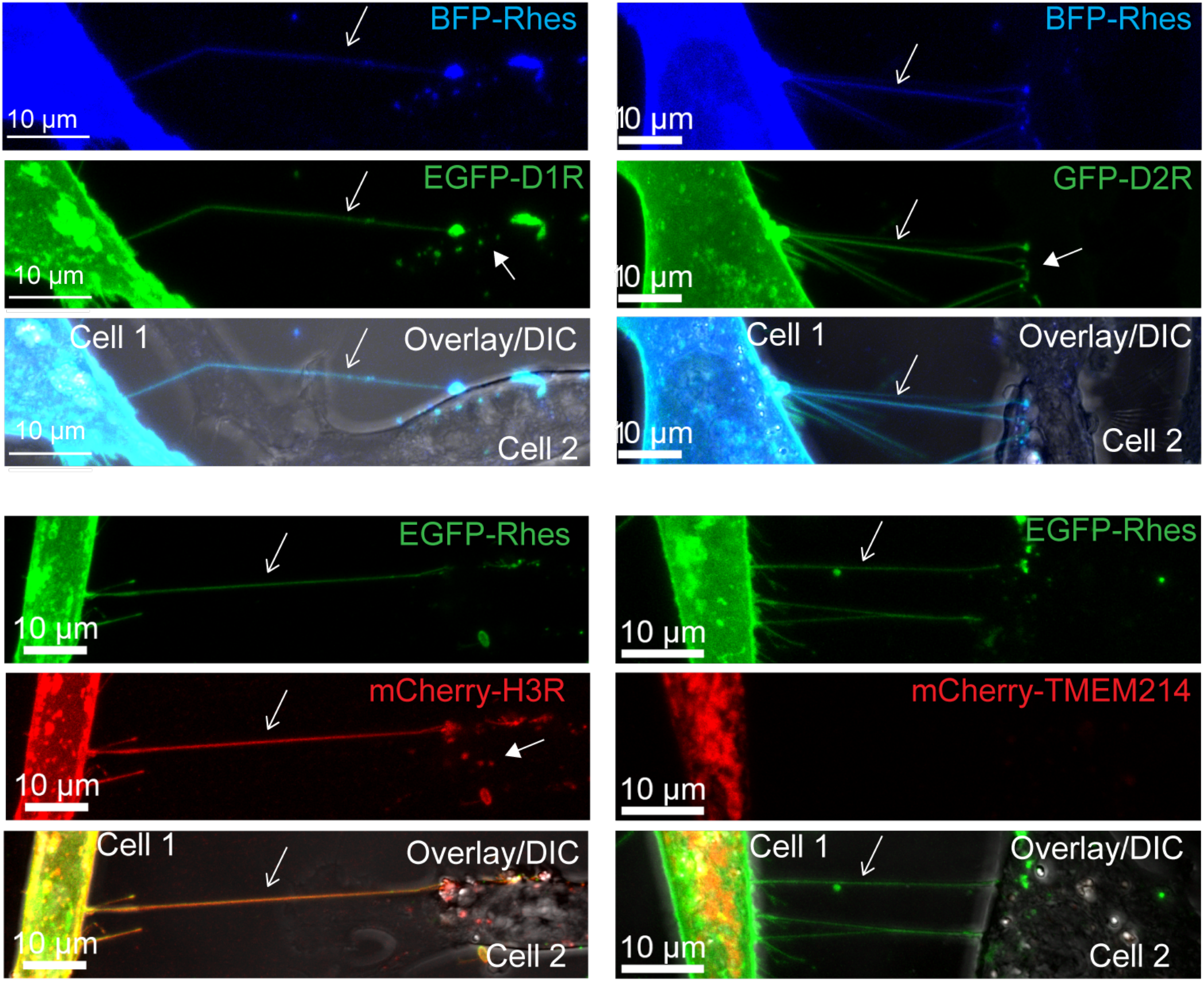
Rhes promotes cargo transport between striatal neuronal cells. Live-cell confocal and differential interference contrast (DIC) images of striatal neuronal cells transfected with indicated reporter constructs. Arrow indicates Rhes tunnel connecting cell 1 to cell 2, and positive for D1R, D2R, H3R but not TMEM214 delivered to cell 2 (arrowhead).

**Supplementary Figure 8.**
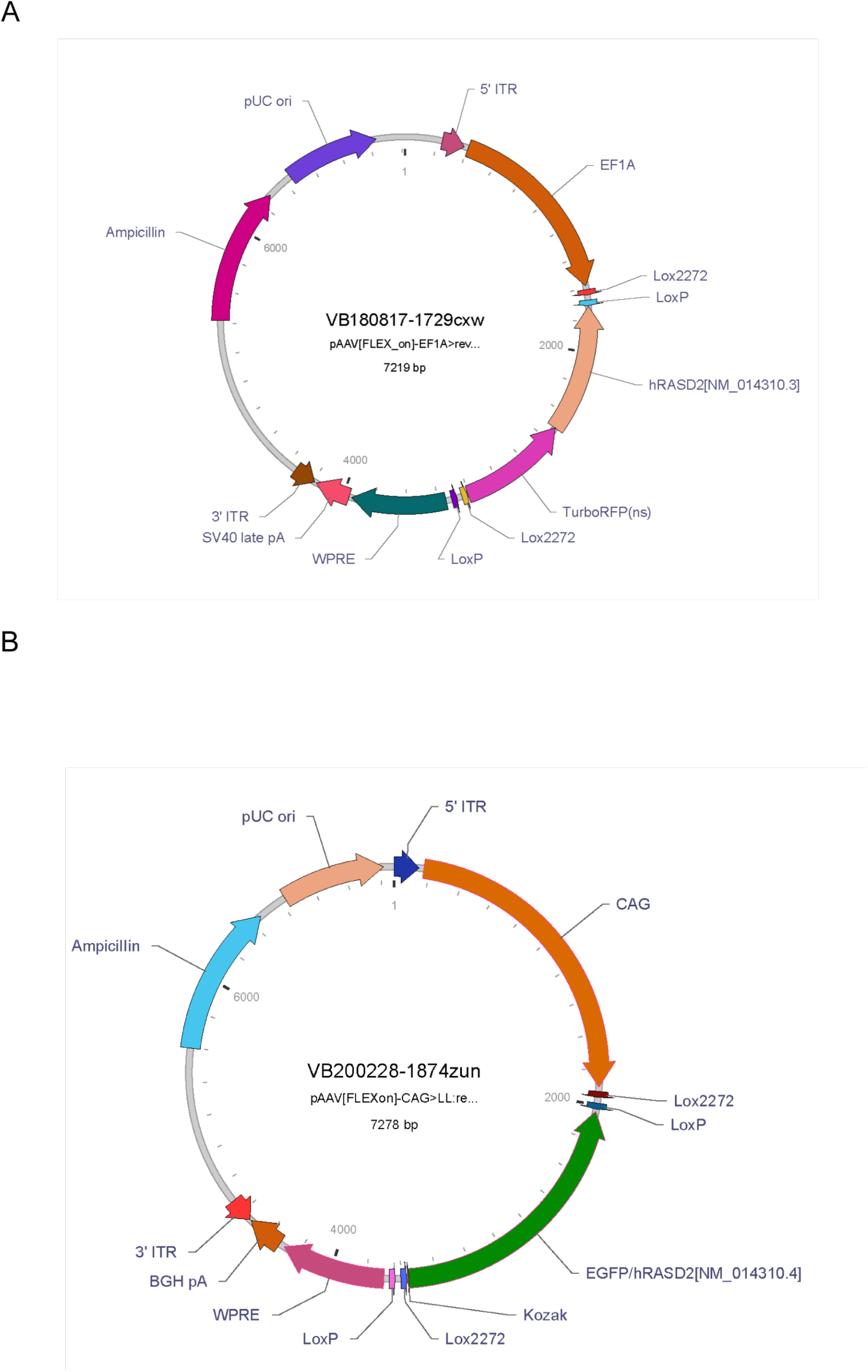
**(A)** Flex Cre-On AAV RFP or AAV RFP-Rhes construct design. **(B)** Flex Cre-On AAV GFP or AAV GFP-Rhes construct design.

